# Temporal chromatin accessibility changes define transcriptional states essential for osteosarcoma metastasis

**DOI:** 10.1101/2022.11.15.516627

**Authors:** W. Dean Pontius, Ellen S. Hong, Zachary J. Faber, Jeremy Gray, Craig Peacock, Ian Bayles, Katreya Lovrenert, Cynthia F. Bartels, Peter C. Scacheri

## Abstract

The metastasis-invasion cascade describes the series of steps required for a cancer cell to successfully spread from its primary tumor and ultimately grow within a secondary organ. Despite metastasis being a dynamic, multistep process, most omics studies to date have focused on comparing primary tumors to the metastatic deposits that define end-stage disease. This static approach means we lack information about the genomic and epigenomic changes that occur during the majority of tumor progression. One particularly understudied phase of tumor progression is metastatic colonization, during which cells must adapt to the new microenvironment of the secondary organ. Through temporal profiling of chromatin accessibility and gene expression *in vivo*, we identify dynamic changes in the epigenome that occur as osteosarcoma tumors form and grow within the lung microenvironment. Furthermore, we show through paired *in vivo* and *in vitro* CRISPR drop-out screens and pharmacological validation that the upstream transcription factors represent a class of metastasis-specific dependency genes. While current models depict lung colonization as a discrete step within the metastatic cascade, our study shows it is a defined trajectory through multiple epigenetic states, revealing new therapeutic opportunities undetectable with standard approaches.

## Main

Over 90% of cancer deaths occur due to metastasis, or the spreading of tumor cells from their original location to other sites within the body ^1^. While the clinical need to target metastasis is clear, the development of anti-metastatic therapies has proved to be difficult for two main reasons. First, large-scale sequencing studies have failed to find genetic aberrations that drive metastatic progression. Unlike tumorigenesis, whose causal mutations are well-characterized in a variety of cancers, metastasis is associated with few recurrent genetic events ^2–4^. Second, the spread of cancer cells to a distal site is a complex, multi-step process known as the metastasis-invasion cascade ^5^. Not only is the mechanism required for initial dissemination distinct from that required for successful growth and survival at the distal site, but each individual stage may be a complex biological process in itself ^6^. For example, the final stage of the metastasis-invasion cascade, known as colonization, encompasses everything from the formation of clinically-undetectable micrometastases to full-blown metastatic disease. During this period of metastasis development, tumor cells are actively adapting to the stresses of a different organ’s cellular millieu in order to ultimately grow in an uncontrolled manner. This transitional middle phase may provide a unique clinical opportunity, yet the underlying biology remains largely unexplored.

Cancer cells accumulate genetic and epigenetic changes during metastatic progression. At the level of the enhancer epigenome, these pro-metastatic changes have been characterized by our lab and others across numerous cancer types ^7–9^. Despite recognition that these changes occur sometime during the evolution of the tumor, when, why, and how they occur remain open questions. Prior studies have relied on profiling and comparing primary tumors to clinically resectable metastases. By limiting comparison to these endpoints, the natural history of cellular states throughout the metastatic cascade is lost. Additionally, while such an approach may be sufficient to capture inherited and somatic genetic mutations that persist in the final output, epigenetic changes are inherently plastic and context-dependent. Thus, they could easily be missed. This leaves the potential for transient intermediate cell states that are undetectable with these static comparisons. Temporal profiling over a defined time course of metastatic colonization is required to investigate this. Genetically engineered mouse models are ideal systems for studying tumor evolution from a single cell, but the stochastic nature of lesion formation makes isolating defined stages of metastasis development challenging. To chart the landscape of epigenetic changes that occur during metastatic colonization, we need a tractable and controlled experimental system with well characterized growth dynamics from metastatic seeding to full colonization. Further, a system amenable to functional perturbation across multiple different human cancer specimens, within the *in vivo* microenvironment, would be ideal.

The lung is the second-most frequent site of cancer metastasis, with twenty-five to thirty percent of all patients with cancer at autopsy having visible lung metastases ^10^. This occurs across many primary tumor types. One of these is osteosarcoma - a particularly aggressive bone cancer that is the second leading cause of cancer deaths in adolescents and young adults ^11^. Despite its favorable prognosis in patients with localized disease, osteosarcoma that has metastasized portends a five-year survival rate below 30 percent - a statistic that has not improved in the last four decades. Here, leveraging osteosarcoma models and functional genomics to study regulatory programs that underlie lung metastasis, we find that colonization is not a single step defined by a single transcriptional program. Instead, it is a trajectory of cell states regulated by distinct factors essential for metastatic progression.

## Results

### Temporal profiling of gene regulation during osteosarcoma metastasis

We performed ATAC-seq and RNA-seq on the metastatic human osteosarcoma cell line MG63.3-GFP grown *in vitro*, as well as harvested from mouse lung at 1 and 22 days after intravenous inoculation (Fig 1A). Although this model of metastasis bypasses dissemination from the primary tumor and intravasation into the circulation, the growth dynamics of subsequent metastases are reproducible and well characterized. Fully formed tumors are present after three weeks, with mice succumbing to metastatic disease at one month. The reproducible nature of this system makes it ideal for studying the regulatory mechanisms underlying colonization. In addition, functional validation is flexible and straightforward due to the ability to perturb gene function in any lung-metastatic human cell line prior to injection. As expected, tumor burden within the lung is visibly different at the two timepoints assessed, with single cancer cells dispersed throughout the lung at day 1, and larger tumors observed at day 22 (Fig 1A). We reasoned that profiling these two timepoints would capture changes associated with both migration/colonization (early steps) and proliferation/outgrowth within the metastatic organ (late steps). Through principal component analysis of the log2 normalized ATAC-seq profiles, we observed clustering of biological replicates from each of the three conditions (Fig 1B). This indicated robust and reproducible epigenomic changes at the time points analyzed. The corresponding analysis of the matched transcriptomes showed similar clustering. These data indicate that the regulatory programs of osteosarcoma cells at the early and late stages of lung metastasis are largely distinct.

**Fig 1:**
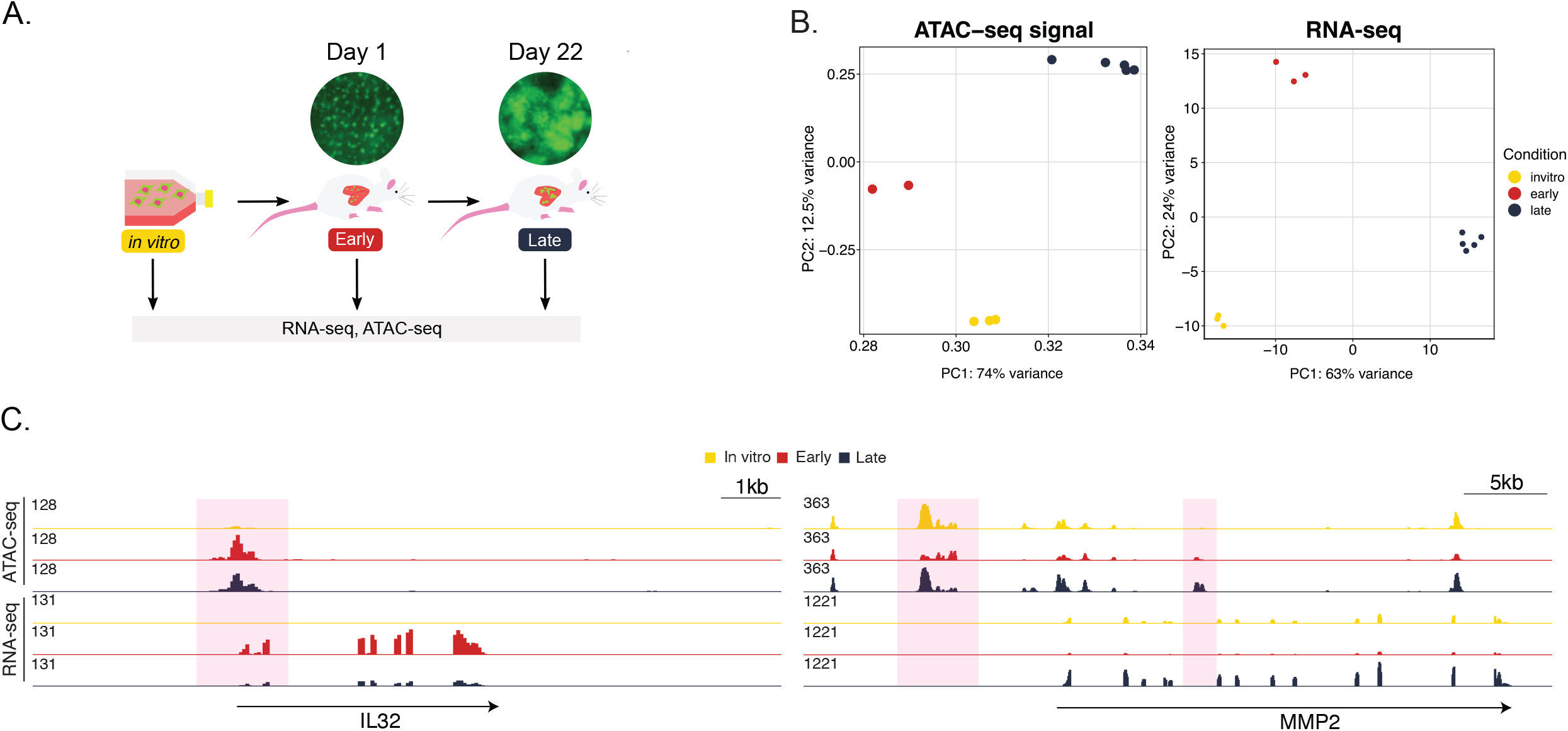
RNA expression and open chromatin are dynamic during osteosarcoma lung metastasis. **A)** Experimental schematic of gene regulation profiling during lung metastasis. Metastatic osteosarcoma cells were injected intravenously. **B)** Principal component analysis of the open chromatin profiles and transcriptomes of cells isolated from three metastasis timepoints. **C)** Representative genome browser screenshots of dynamic accessible regions and corresponding transcripts.

Regions that change in accessibility during metastatic progression are distributed throughout the genome, with some changes occurring at gene promoters, and others occurring at intergenic or intronic regions (Fig 1C). Many of the associated genes have been investigated in the context of osteosarcoma metastasis, but underlying mechanisms contributing to their aberrant expression remain unclear. For example, the gene *IL32*, which encodes a cytokine shown to promote osteosarcoma cell invasion and motility, displays an increase in chromatin accessibility at its promoter primarily at the early time point ^12^ (Fig 1C). This increase in accessibility is reflected in the gene’s expression. In addition, the collagenase encoded by *MMP2* has been associated with osteosarcoma pulmonary metastasis ^13^. Our data demonstrate that *MMP2* expression increases at the late *in vivo* timepoint, and is associated with an increase in chromatin accessibility at a putative intronic enhancer element (Fig 1C).

To determine if these findings generalize to other specimens, we repeated the experiment using another metastatic osteosarcoma cell line model: 143b-HOS-GFP (Supp. Fig. 1). This revealed that dynamic changes in chromatin accessibility also occur in 143b-HOS-GFP, with the three different conditions again clustering separately at both the chromatin and RNA levels (Supp. Fig. 1A-B). While the exact regions that changed in accessibility varied between the two cell lines, there was a degree of overlap (Supp. Fig. 1C). For example, the promoter of *IL32* again displayed an early-specific increase in accessibility (Supp. Fig. 1D). This implies common biology underlies the different stages of osteosarcoma metastasis despite genetic and epigenetic variation, and that these programs are regulated by reproducible changes in the epigenomes of metastasizing cells.

### Dynamic shifts in chromatin accessibility during osteosarcoma progression correlate with temporally-distinct metastatic programs

We used k-means clustering to partition the dynamic epigenome of MG63.3-GFP cells based on how each region changes in chromatin accessibility over the metastasis time-course. We found eight robust clusters (Fig 2A-B). The first four clusters contained regions more accessible at one or more of the *in vivo* time points, compared to *in vitro*. We thus annotated them as early (cluster 1), pan-*in vivo* (clusters 2 & 3), and late (cluster 4), and will refer to them as such throughout the rest of the text. In addition to the clusters of peaks gained *in vivo*, we also identified clusters that were lost throughout metastasis (cluster 6), or at a specific time during metastatic progression. Cluster 5, for example, contains regions that transiently decrease in accessibility early, while cluster 8 is composed of regions that are relatively inaccessible late.

**Fig 2:**
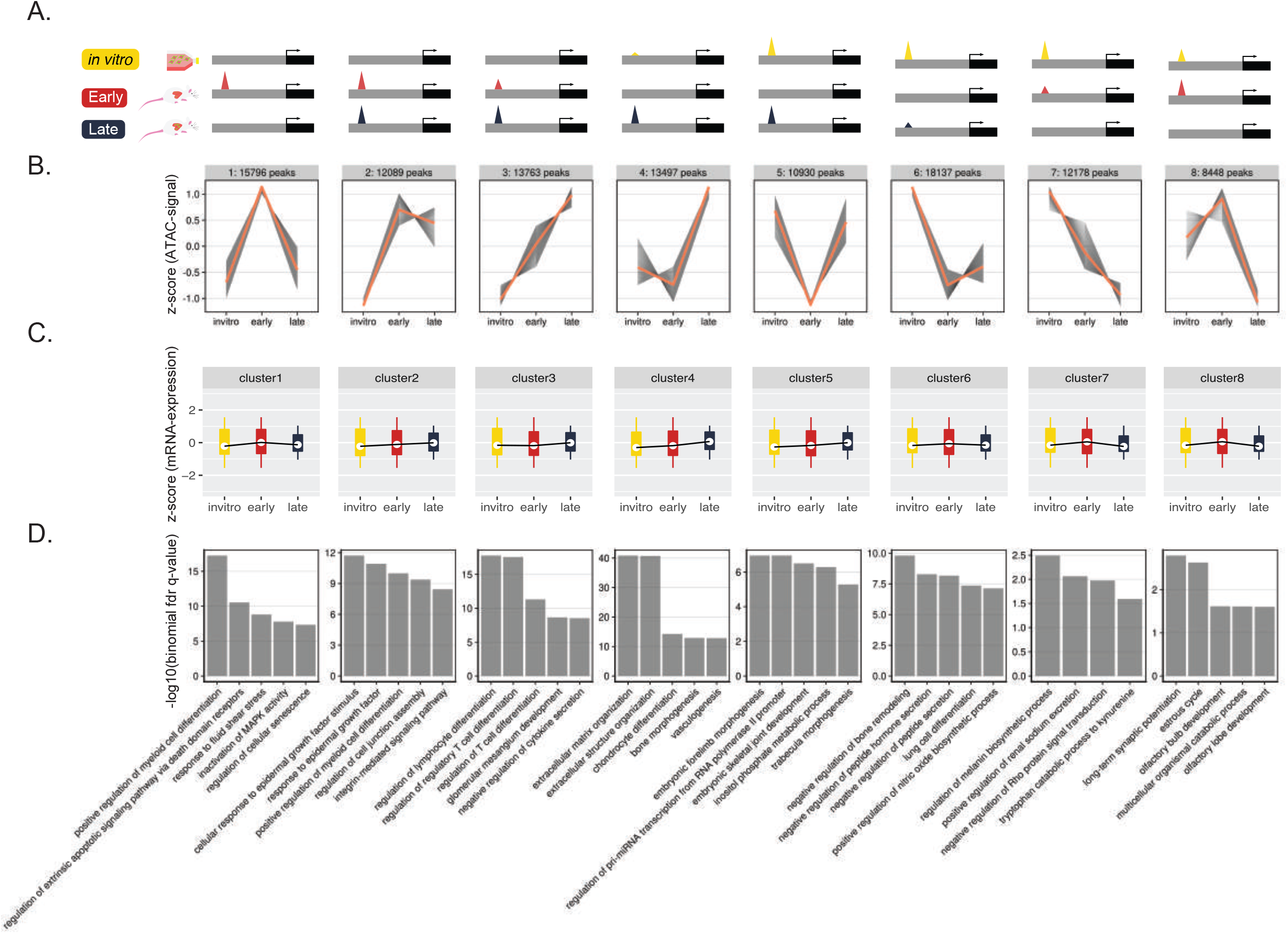
Dynamic shifts in chromatin accessibility during osteosarcoma progression regulate temporally-distinct metastatic programs. **A)** Diagram illustrating representative genome-browser views for peaks within each dynamic cluster. **B)** K-means clustering partitions the “universe” of open chromatin based on their accessibility dynamics over time. Grey lines represent the dynamics of an individual peak, while the orange line represents the mean change for all peaks within a given cluster. **C)** Z-scored mRNA expression for genes associated with peaks within each cluster. **D)** Peak ontology based on GREAT for peaks within each cluster. Top significant terms are shown.

Positive cis-regulatory elements like enhancers and promoters lie within regions of accessible chromatin ^14^. However, accessible chromatin also houses insulator elements that help constrain three-dimensional genomic interactions within confined regions, and are bound by structural transcription factors such as CTCF ^14^. If the regions we identified are acting predominantly as functional enhancers or promoters, we would expect the expression of their target genes to mirror the changes in accessibility. Analysis of the expression of target genes predicted by the Genomics Regions Enrichment of Annotations Tool (GREAT) revealed this was indeed the case for the first 4 clusters (Fig 2C) ^15^. We found that genes associated with the early cluster (1) displayed the highest level of expression at the early timepoint. Similarly, genes paired with the pan-*in vivo* clusters (2 & 3) showed higher expression in both *in vivo* conditions when compared to *in vitro*. This same phenomenon held true for the late cluster (4), whose target genes were expressed highest at the late timepoint. However, correlation between expression and accessibility was weak for the other clusters. This implies the accessible regions in clusters 5 through 8 may include architectural or negative regulatory elements such as insulators/silencers that dominate the control of associated genes.

We used GREAT to identify biological processes associated with the three *in vivo* gain clusters (early, pan-*in vivo*, and late; Clusters 1 – 4) (Fig 2D) ^15^. Remarkably, biological terms for each cluster were quite distinct, and included processes with clear ties to metastasis. This suggests that the clusters regulate gene programs corresponding to distinct stresses faced throughout metastasis. The early regions (cluster 1) regulate genes associated with fluid shear stress, a likely response to the mechanical forces present during migration through the circulation, and extravasation into the secondary organ. Additionally, this same cluster seems to be responsible for regulating apoptosis and senescence two pathways critical for survival of micrometastases in the stressful new microenvironment of the lung ^16,17^. The pan *in vivo* regions (clusters 2 & 3) are enriched for terms involving extracellular stimuli, such as response to growth factors and cell-to-cell interactions. These are biological processes required for osteosarcoma cells to interact with the new cell types that make up the milieu of the metastatic microenvironment. Lastly, the late regions (cluster 4) associate with pathways important for the growth of larger metastatic lesions. These pathways include remodeling of the local extracellular matrix, activation of bone development programs important for osteosarcoma growth, and activation of vasculogenic programs required to support the increased nutrient and waste transport needs of larger tumors. In the 143b-HOS-GFP cell line, we see partial overlap of the significantly enriched terms for each cluster (Supp. Fig. 2A-B). This indicates that even in tumors with genetically diverse backgrounds, common transcriptional programs are required to successfully navigate the different stages of lung metastasis represented in our model.

### Distinct transcription factors regulate temporal chromatin clusters

Transcription factors mediate the regulation of genes through context-specific interactions with enhancers and promoters ^18^. To identify which transcription factors are controlling the temporally-distinct changes in chromatin accessibility, we analyzed the DNA sequence within each cluster of accessible regions to find differentially enriched motifs (Fig 3A). This analysis revealed unique sets of TFs predicted to bind to the various accessible chromatin clusters. While the early regions were highly enriched for FOX motifs and NFkB motifs, the pan *in vivo* regions showed specific enrichment of motifs for a variety of KLF factors. Interestingly, the late peaks were enriched for known mesenchymal factors like TWIST1 and SOX9/10, which have been previously shown to play a role in bone and cartilage development ^19,20^. Of note, regions of the genome that lose accessibility early in metastasis (cluster 5) are highly enriched for CTCF motifs. Since CTCF is a structural factor involved in securing three-dimensional chromatin interactions, the loss of CTCF-enriched accessible sites may represent a loss of insulation and reorganization of chromatin topology ^21^. To determine if these transcription factors were regulating the equivalent chromatin dynamics in other contexts, we performed the same analysis on the accessible chromatin clusters characterized in 143b-HOS-GFP (Supp. Fig. 2C). NFkB family motifs were again enriched in the cluster 1 (early), while SOX9 and TWIST1 motifs were enriched for cluster 4 (late), demonstrating generalizability of our findings.

**Fig 3:**
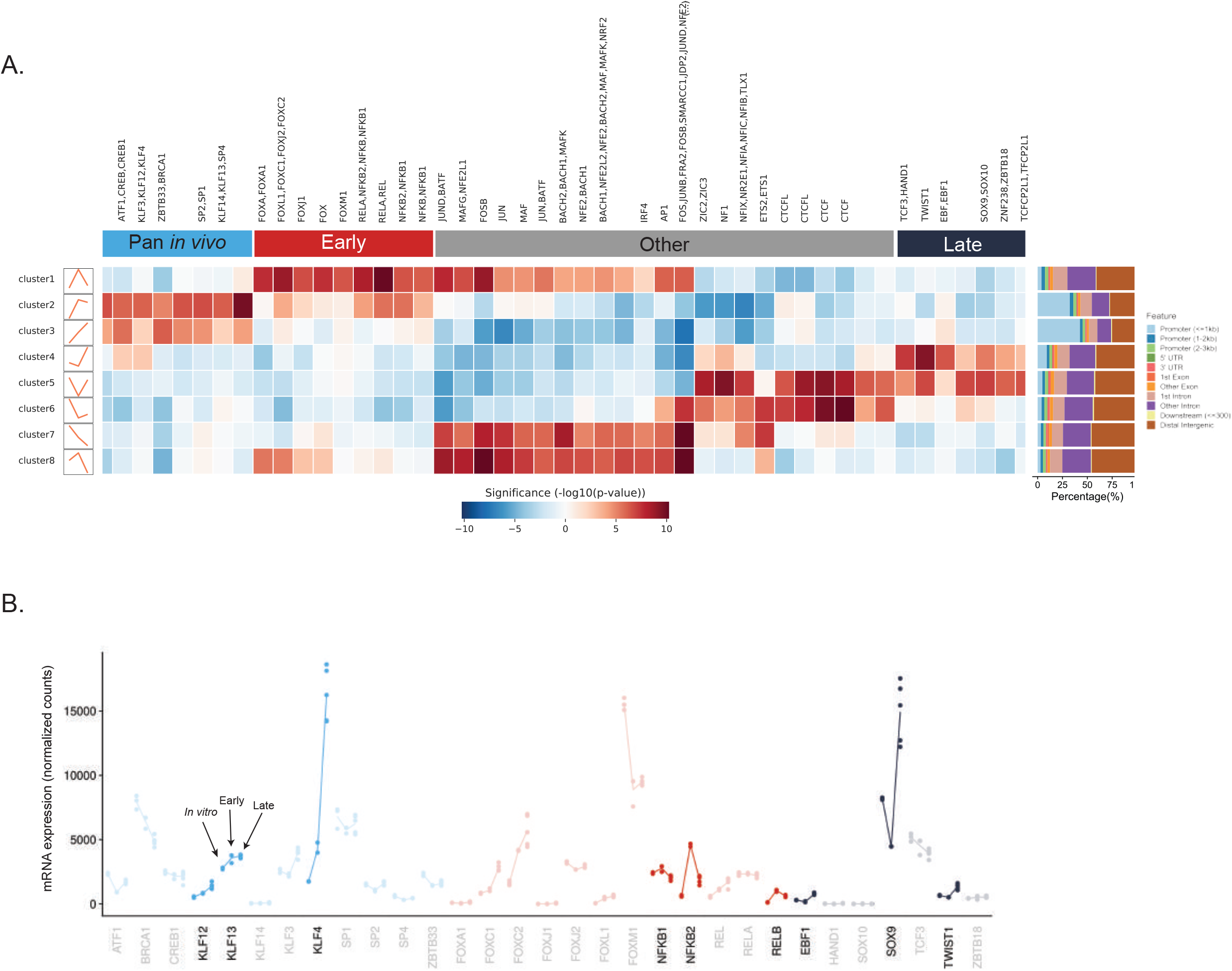
Putative regulators of dynamic-clusters are identified through motif-mining and gene expression changes. **A)** Transcription factor motif enrichment heatmap displaying differentially enriched motifs for each dynamic cluster with genomic annotation of accessible regions contained within each cluster. Blank heatmap columns correspond to *de novo* motifs. **B)** Gene expression dynamics of motif-enriched transcription factors during metastatic progression. Highlighted genes show expression changes that mirror the cluster’s accessibility dynamics.

To further pare down the list of putative regulators, we used the timecourse RNA-seq data to identify TFs whose expression dynamics mirrored the accessibility of the peaks within their respective cluster (Fig 3B). Out of the factors enriched in the early-specific cluster, *NFKB2, NFKB1*, and *RELB* stood out as having an early-specific increase in expression, indicating a potential for differential activity at this time point. Excitingly, two of these genes, *NFKB2* and *RELB*, converge at the pathway level, as both are components of non-canonical NFkB signaling - a known driver of metastasis in other chromosomally unstable cancers ^22^. For the putative pan-*in vivo* cluster regulators, *KLF4, KLF12*, and *KLF13* all show an increase in expression at both *in vivo* timepoints compared to the same cells profiled *in vitro*. Lastly, we identified *SOX9, EBF1*, and *TWIST1* as likely late-cluster regulators through this same approach. Many of these timepoint-specific patterns of TF gene expression were also observed in another metastatic osteosarcoma cell line (MNNG-HOS) when the cells were grown within lung explants (Supp. Fig 3). This supports the idea that a common set of regulators are responsible for the microenvironment-induced epigenomic reprogramming events across osteosarcomas.

While these accessibility dynamics could be due to active chromatin changes *in vivo*, they could also result from selection of subpopulations of cells that already possess the observed profiles *in vitro* (Supp. Fig 4A). We performed single cell ATAC-seq on MG63.3-GFP cells grown *in vitro* to investigate these two possibilities (Supp. Fig 4B). Using the above TFs as markers for the temporal accessible chromatin landscapes, we projected either 1) their motif enrichment or 2) their promoter accessibility onto the scATAC-seq UMAP space (Supp. Fig 4C). This showed the unbiased clusters identified *in vitro* were not defined by temporal TF activity, indicating the dynamic peaks are likely a result of microenvironment-dependent reprogramming events, and not subclonal population shifts.

### The middle phase of lung colonization harbors condition-specific dependencies

While our data and integrated analyses define a sequence of epigenomic reprogramming events that occur during metastatic progression, it is unclear whether these changes are necessary for metastasis to occur. We hypothesized if these changes were indeed driving metastasis, the upstream regulators would be essential. *In vitro* genome-scale CRISPR dropout screens have emerged in the past decade as a powerful way to assess gene dependency in a high-throughput manner. However, *in vivo*-specific biology is not captured with traditional workflows that solely use *in vivo* models to validate *in vitro* dependencies. In addition, screening coverage, and thus data quality, are limited by the number of cells that initially engraft within the lung. This makes genome-scale dropout screens in models of lung metastasis experimentally infeasible. To overcome these barriers, we designed a single guide RNA (sgRNA) library to target 78 TFs (along with 11 positive controls and 25 non-targeting negative control sgRNAs), and performed parallel experiments *in vitro* and in an *in vivo* model of lung metastasis (Fig 4A). By combining 4 mice per replicate, we were able to achieve a screening coverage of 400x *in vivo*. Both screens were performed using the MG63.3-GFP-Cas9i cell line previously described ^23^.

**Fig 4:**
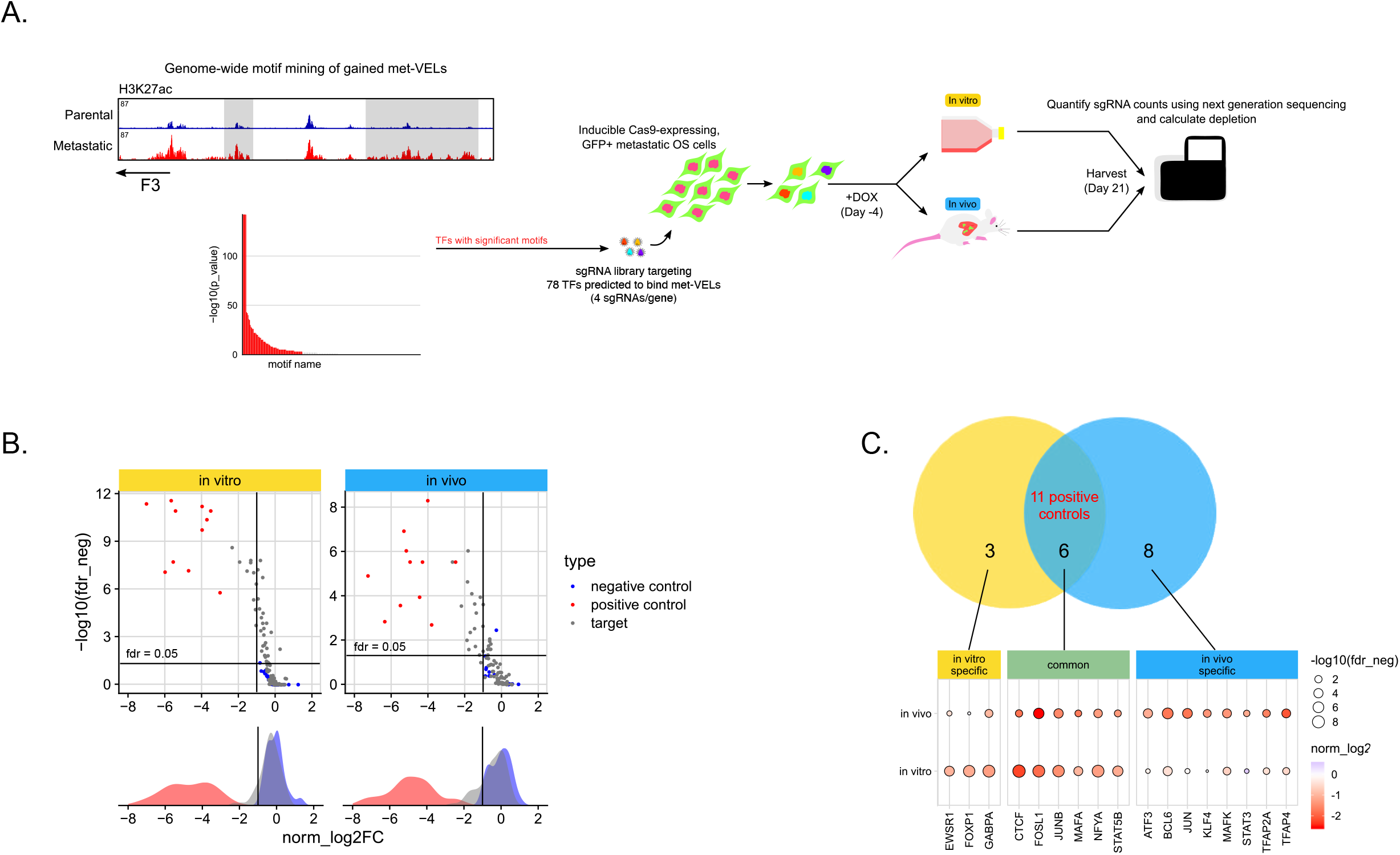
Targeted CRISPR screen reveals condition-specific transcription factor dependencies in metastatic osteosarcoma. **A)** Experimental workflow for *in vitro* and *in vivo* CRISPR screens in a metastatic osteosarcoma cell line. **B)** One-sided volca- no plots and marginal density plots displaying distribution of gene targets included in the screens. Gene-level statistics are shown. **C)** Set of genes called as hits in each screen. The cutoff used for calling hits was an fdr neg < 0.05 and log(fold-change) < -1.

We confirmed significant depletion of sgRNAs targeting all 11 positive controls in both arms of the screen. In addition, the non-target negative controls did not affect cell fitness, collectively demonstrating the quality of both screens (Fig 4B). In total, the knockout of 17 TFs were shown to reduce cell growth both *in vitro* and within the lung microenvironment. Excitingly, 8 TFs were classified as metastasis-specific dependency genes, with fewer classified as *in vitro*-specific (3 genes) or common dependencies (6 genes other than the positive controls) (Fig 4C). The 8 metastasis-specific dependency genes included multiple TFs from the AP-1 factor family, which has previously been implicated in osteosarcoma metastasis ^7,24^. Besides these, we found *KLF4, STAT3, BCL6, TFAP2A*, and *TFAP4* as potentially novel drivers of osteosarcoma metastasis. To further investigate the context-specificity of our genes of interest, we probed publicly available whole genome CRISPR screen data in 990 cell lines from the Dependency Map (DepMap) database ^25,26^. While *TFAP4* was an essential gene in 48% of the cell lines and *BCL6* was a lymphoma-specific essential gene, the other 6 *in vivo* hits showed little evidence of depletion across all cancer types, including 9 osteosarcoma cell lines (Supp. Fig. 5). This finding further highlights the importance of screening directly in the metastatic microenvironment, and that our *in vivo* hits are selectively important for lung metastasis, and not other osteosarcoma growth contexts.

### Genetic and pharmacological inhibition of pro-metastatic transcription factors prevents lung metastasis

We reasoned that if these TFs were indeed metastasis-specific dependencies then inhibiting them should impair lung metastasis but spare *in vitro* growth. We first sought to show that these factors demonstrated context-specific importance outside of the screening setting. Since *KLF4* was not only an *in vivo*-specific hit from our screen, but also a likely mediator of the *in vivo*-specific chromatin changes observed, we chose to start with this factor. Using the top sgRNA from our screening library, we knocked out *KLF4* in a pooled format in MG63.3-GFP-Cas9i (Fig 5A). As a control, we transduced the same cell line with a non-targeting sgRNA. We observed that knocking out *KLF4* did not decrease the viability of metastatic osteosarcoma cells grown over the course of 7 days *in vitro* (Fig 5B). In line with these findings, knocking out *KLF4* also had no impact on the *in vitro* enhancer landscape of MG63.3-GFP cells (r = 0.98) (Supp. Fig. 6A-B). Using CRISPResso2, we determined the majority (85%) of ChIP-seq reads mapped to the *KLF4* locus in our knockout cells contained edits predicted to disrupt KLF4 expression (Supp. Fig. 6C) ^27^. This was not the case for the *KLF4* wild-type ChIP-seq. This confirmed the similarity of the enhancer landscapes between the two cell lines was not simply due to outgrowth of *KLF4* wild-type cells within our pool.

**Fig 5:**
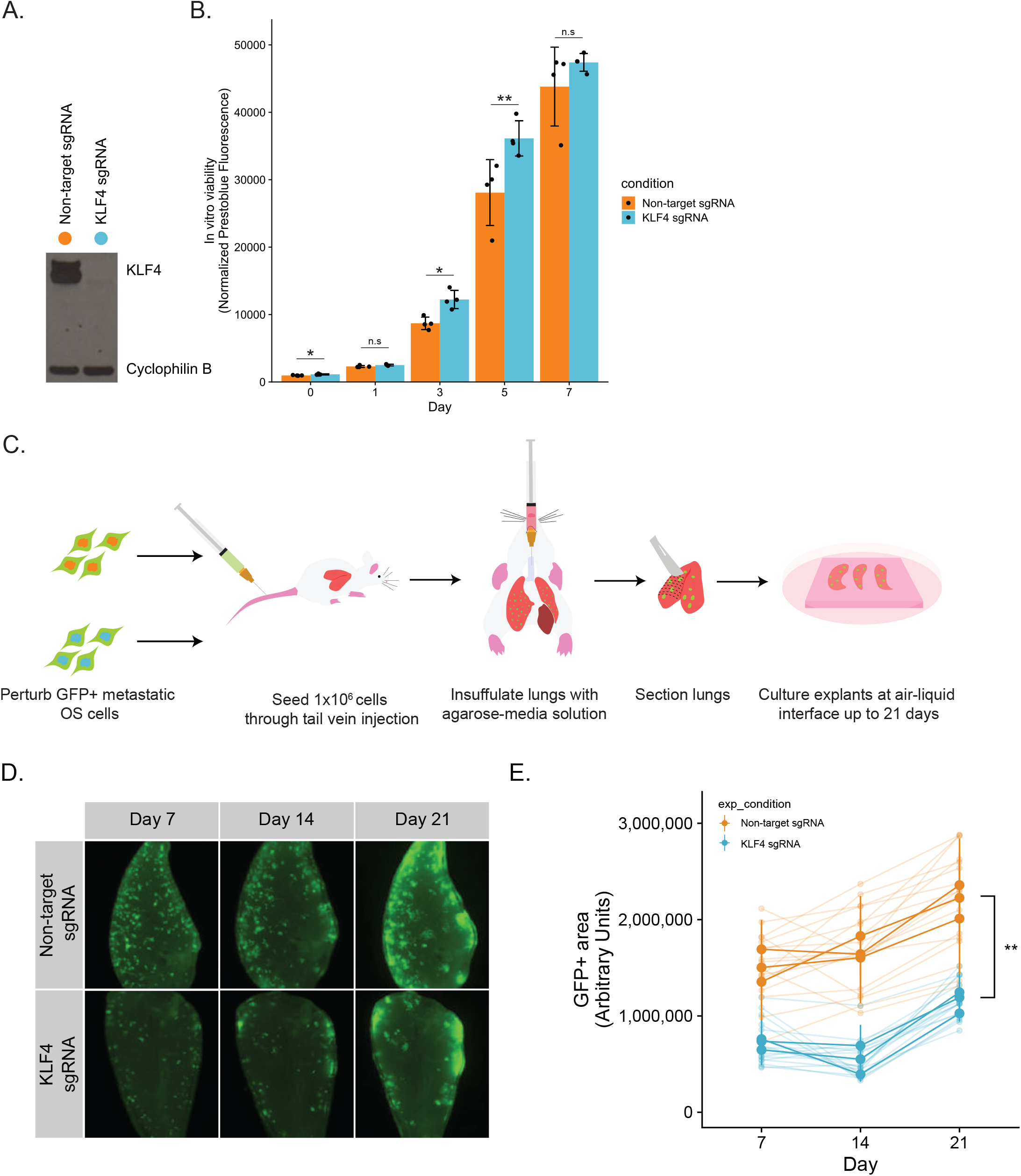
Kruppel-like factor 4 (KLF4) is a *bona fide* metastasis-specific dependency. **A)** Western blot showing KLF4 knockout in a metastatic osteosarcoma cell line compared to non-target control transduced cells. **B)** *In vitro* cell viability assay using presto-blue reagent. N = 4 per cell line. **C)** Illustration of the workflow for the *ex vivo* pulmonary metastasis assay (PuMA). **D)** Longitudinal imaging of representative lung sections for each cell line. **E)** Quantification of metastatic burden in the PuMA assay using GFP+ area.

In order to study the effect of *KLF4* knockout on metastatic capability, we used an *ex vivo* Pulmonary Metastasis Assay (PuMA) - a secondary model of lung metastasis ^23,28^. In this system, metastatic osteosarcoma cells are injected into the lungs of mice through the tail vein (Fig 5C). The mice are immediately euthanized, after which lung slices are grown at an air liquid interface *ex vivo*. By monitoring the tumor burden by GFP-positive area within each lung slice, we saw that knocking out *KLF4* decreased the ability of the cells to grow within the context of the metastatic microenvironment. At every time point imaged, the level of tumor burden was significantly lower in the *KLF4* knockout cell line than the non-target control line (Fig 5D-E). While tumor burden did increase over time for both conditions, this could be representative of cells within the pool of *KLF4* knockout cells that escaped editing of the *KLF4* locus. However, we cannot rule out the possibility that KLF4 is only partially essential for lung metastasis, or that metastasizing cells can circumvent KLF4 dependence through usage of alternative transcriptional machinery.

Despite the clear importance of KLF4 in promoting metastasis in our models, therapeutic inhibitors of this factor have not yet been developed. However, other *in vivo* dependencies from our screen, such as STAT3, have chemical probes that can be used to impair their function. We thus sought to determine if a STAT3 inhibitor could selectively target osteosarcoma lung metastasis.

Using the PuMA system, we found that the STAT3 inhibitor TPCA-1 prevented *ex vivo* lung metastasis at 5 uM and 1 uM over the course of 21 days (Fig 6A-B). Confirming our genetic findings that *STAT3* is an *in vivo*-specific dependency, the same cells were unaffected by TPCA-1 when grown outside of the lung microenvironment (Fig 6C). These data demonstrate that our approach identified druggable metastasis dependency genes that may serve as therapeutic targets for treating metastatic osteosarcoma.

**Fig 6:**
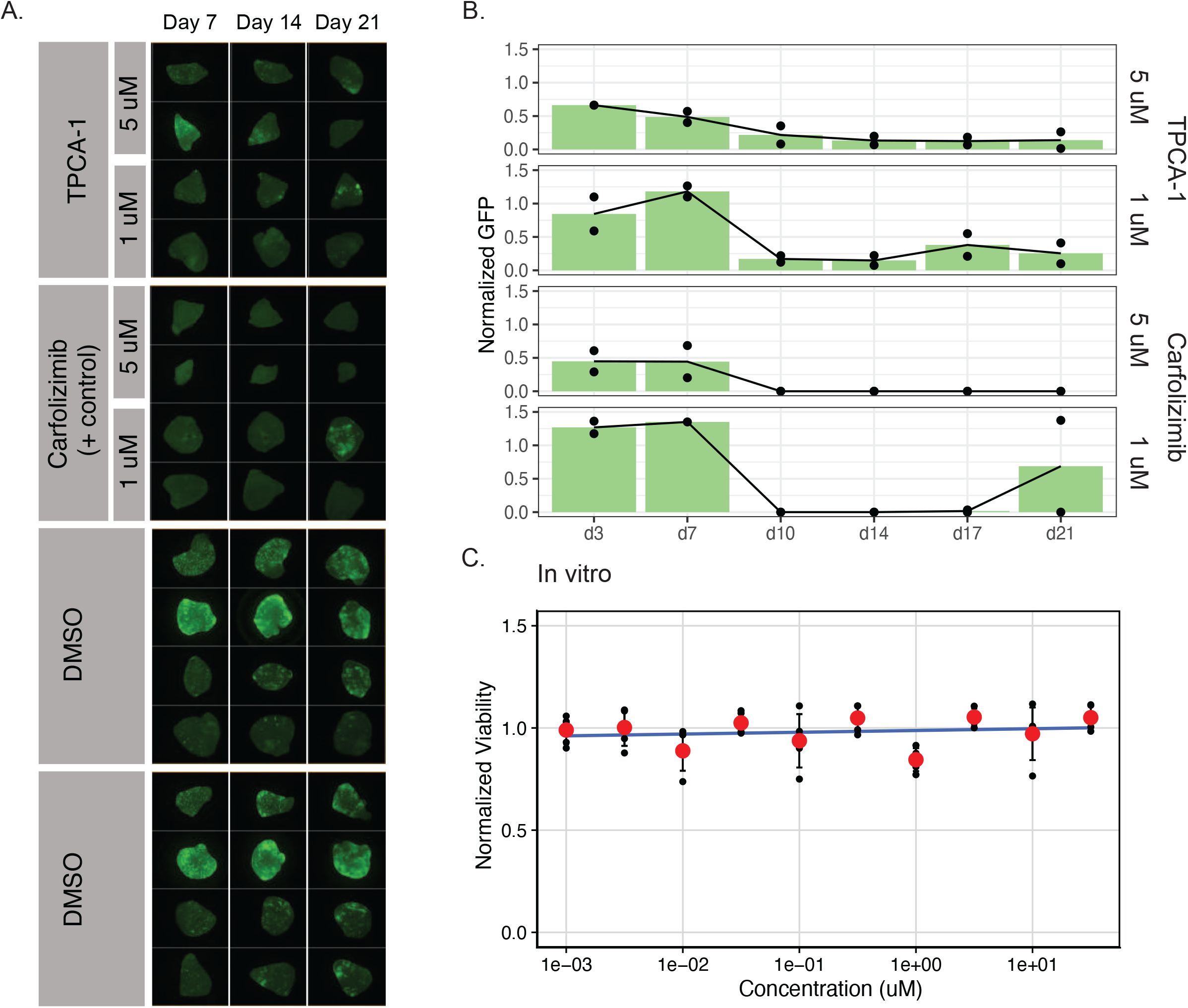
STAT3 inhibition with TPCA-1 attenuates metastasis without *in vitro* cell viability effects. **A)** Longitudinal imaging of PuMA lung sections treated with TPCA-1, Carfolizimib (positive control), or DMSO (negative control). **B)** Quantification of metastatic burden for each condition based on GFP signal. Treatment conditions were normalized to DMSO control sections from the same mouse to account for intermouse variability. **C)** *In vitro* dose response curve for MG63.3 cells treated with TPCA-1. PrestoBlue reagent was used to measure cell viability.

### Metastasis dependency transcription factors represent a transcriptional addiction

While some of the pro-metastasis TFs are targetable with specific inhibitors, not all TFs are able to be directly targeted in this manner. However, many of the metastasis-specific hits from our *in vivo* screen are upregulated at the mRNA level within the lung microenvironment in 2 cell lines (Fig 7A-B). This indicates that metastatic osteosarcoma cells may depend on the increased expression of these TFs for successful lung colonization. In other words, the dependence on this specific set of transcription factors may represent a metastasis-specific transcriptional addiction. We wondered whether this increase in expression of pro-metastasis TFs could be a potential vulnerability for metastatic osteosarcoma cells.

**Fig 7:**
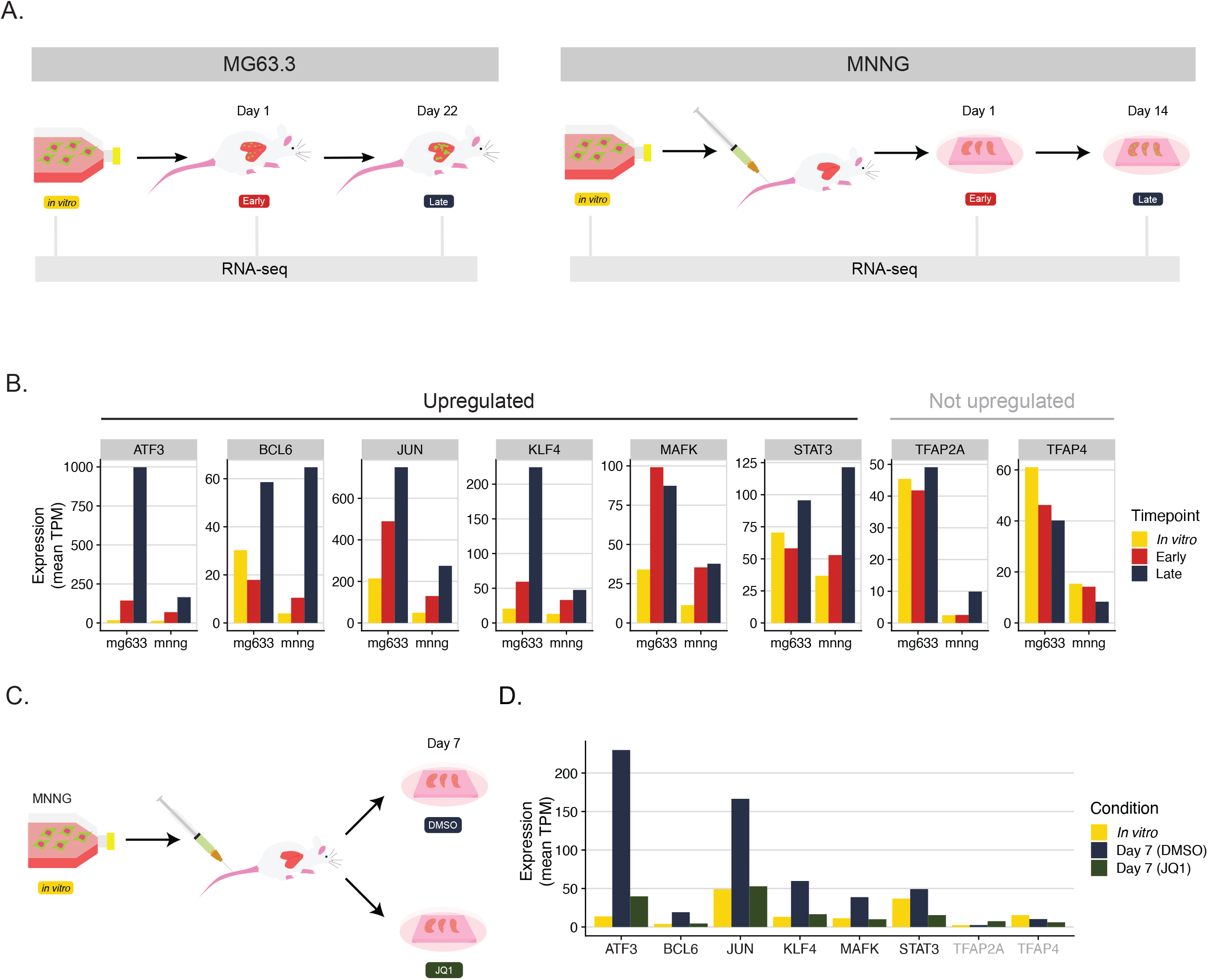
Transcriptional inhibition with JQ1 prevents lung metastasis and abrogates the increased expression of pro-metastatic transcription factors within the lung microenvironment. **A)** Schematic comparing RNA-seq profiling experiments of MG63.3 and MNNG cell lines. MG63.3 early and late conditions were isolated from a fully *in vivo* model of metastasis, whereas MNNG cells were isolated from the ex vivo PuMA model. **B)** Barplots demonstrating RNA expression dynamics of *in vivo* CRISPR screen hits in both MG63.3 and MNNG cells. **C)** Schematic illustration of experiment assessing effect of JQ1 on MNNG gene expression during lung metastasis. **D)** RNA expression of *in vivo* CRISPR screen hits in the MNNG cell line, with and without JQ1 treatment.

Experimental probes that target the transcriptional machinery such as JQ1 and THZ1 have been used to determine transcriptional addictions in cancer cells ^29–31^. Treatment of the metastatic osteosarcoma cell line MNNG-HOS with JQ1 has been shown to block tumor growth within the lung microenvironment ^7^. To determine if the anti-metastasis property of JQ1 correlated with downregulation of any of the pro-metastatic transcription factors identified, we reanalyzed data profiling the transcriptome of MNNG-HOS grown in the PuMA system with and without drug. Strikingly, 6 of the 8 *in vivo* hits from MG63.3 showed an increase in expression when MNNG-HOS cells were grown within the context of the lung microenvironment (Fig 7B). The increase in expression of all 6 of these factors was attenuated by exposure to JQ1 (Fig 7C-D). These data indicate that transcription factor dependencies identified in one cell line could be more broadly relevant to human osteosarcoma, and transcriptional inhibition may serve as a general therapeutic strategy.

## Discussion

Despite metastasis being a devastating clinical problem, our understanding of the epigenetic mechanisms associated with the process have been limited to the biological “bookends”. While mutational drivers of cancer processes are preserved during tumor progression, the plasticity of the epigenome means epigenetic drivers can be transient. This can make them invisible with traditional end point comparisons. The past focus on comparing primary tumors to late-stage metastatic disease has limited our understanding of the complex phase of lung colonization, which remains a black box. Here we show this critical period of osteosarcoma metastasis is composed of a continuum of changes that occur as cancer cells interact with their new microenvironment.

Waddington’s epigenetic landscape for embryonic development depicts a series of ridges and valleys that represent paths a pluripotent cell can take to differentiate into a variety of somatic cell types. Along each of these paths, the differentiating cell encounters branch points that end in different tissues, with the apex of this branch point representing intermediate cell states. In a similar way, we envision a metastasizing cell as rolling down an analogous epigenetic landscape. Where fully differentiated normal tissue is present at the bottom of Waddington’s landscape, our model places metastatic lesions indicative of end stage disease. Our work defines the epigenetic path an osteosarcoma cell takes to colonize the lung, identifying multiple intermediate cell states along the way.

Importantly, we found these specific epigenetic reprogramming events present vulnerabilities that can be targeted at multiple distinct levels of gene regulation. Upstream transcription factors driving this reprogramming can play temporally-specific roles in the process of lung colonization. However other factors like the master transcription factor KLF4 are ubiquitously important. We show through an *in vivo* functional genomics approach and subsequent validation experiments that KLF4 is indeed required for successful outgrowth in a murine pulmonary metastasis xenograft model. This same factor was totally dispensable for the same cells to grow *in vitro*, emphasizing its context-specific importance. More work is required to determine correlates of KLF4 dependence, such as patient sex, tumor location, or specific genomic/epigenomic alterations (i.e. amplification of *MYC*, mutations in *TP53*, specific variant enhancers, etc).

In addition to KLF4, we identify seven other transcription factors that are essential for lung metastasis, but unimportant for *in vitro* proliferation. One of these factors, STAT3, can be directly targeted in an experimental setting using the drug TPCA-1. However, more broadly, the general addiction of metastasizing cancer cells to the expression of TF dependency genes is indirectly targetable through transcriptional inhibition with JQ1. This provides further evidence that multiple different approaches to targeting the transcriptional axis show promise as therapeutic strategies in metastatic osteosarcoma.

Although we focus mostly on pan *in vivo* dependencies, another exciting finding from our work is that the different stages of metastasis may be targeted by inhibiting the specific transcriptional machinery required for each phase of lung colonization (Fig 8). This indicates targeting the various clinical manifestations of the disease will likely require different drugs. For example, targeting early regulators will likely be optimal for preventing early phases of colonization, whereas late regulators may be better targets for more established metastatic disease.

**Fig 8:**
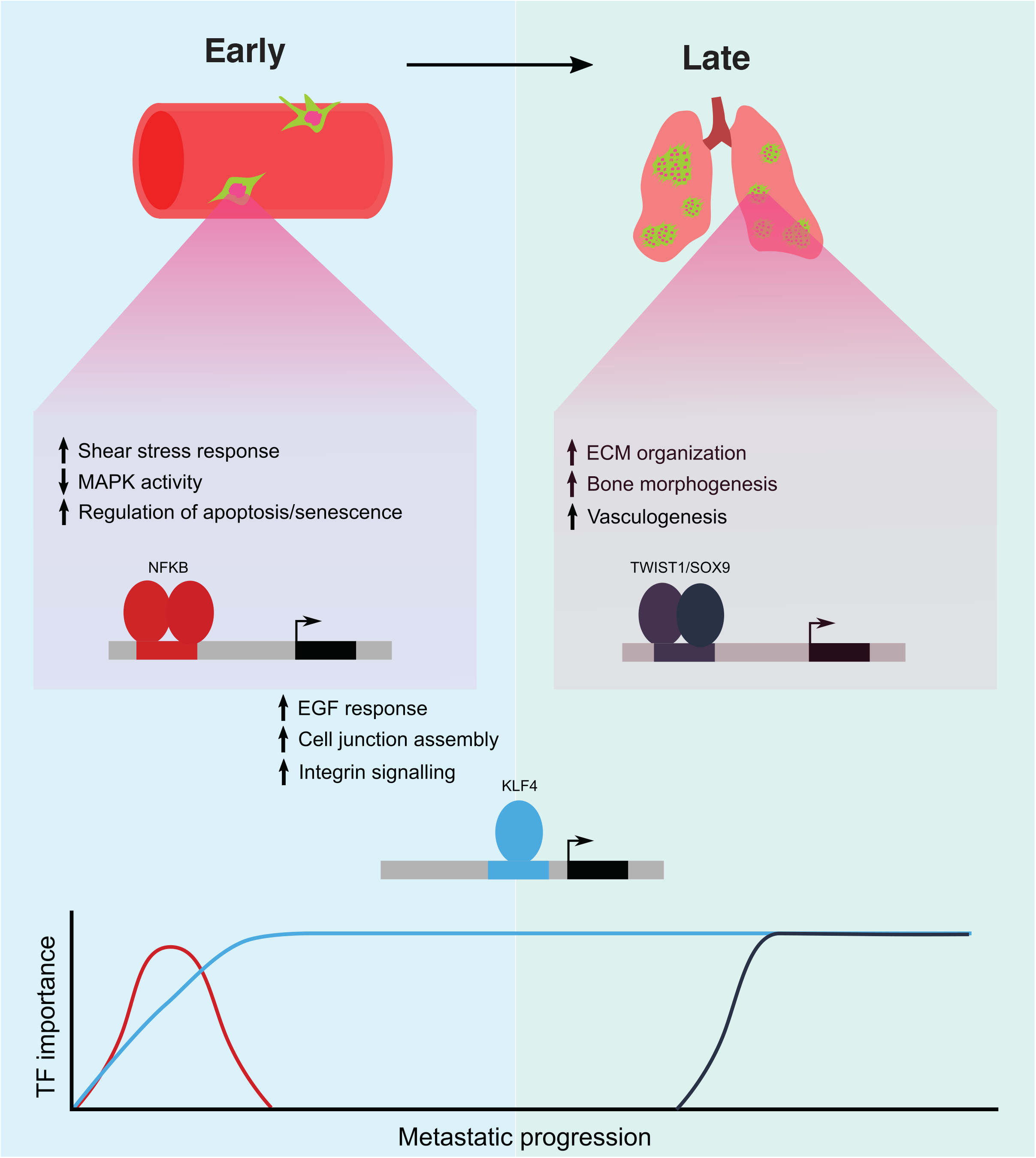
Model for transcription-factor driven lung metastasis in osteosarcoma.

Our work also provides novel insights into the basic biology underlying osteosarcoma lung metastasis. Both SOX9 and KLF4 have been shown to play key roles in normal bone development. KLF4 inhibits osteoblast differentiation to regulate bone homeostasis, while SOX9 serves as a master regulator of chondrocytes - an intermediate cell type during osteoblastogenesis ^32,33^. Thus, our data suggest metastatic competence may depend on osteosarcoma’s ability to transiently dedifferentiate from an osteoblast-like state. This is in line with findings in prostate cancer showing early developmental programs are important for metastasis ^34^.

In totality, we develop a strategy combining metastasis time point experiments and *in vivo* functional genomics to elaborate on seminal work describing the importance of the epigenome to the metastatic phenotype across multiple cancers ^7–9,35^. While other studies focused on end point comparisons, we are the first to show distinct changes at the level of chromatin occur multiple times throughout lung colonization. Furthermore, these temporally resolved changes point in the direction of a host of novel metastasis-dependency genes that lie throughout the dynamic epigenomic landscape of metastasizing cells. Experimentally, this study demonstrates the importance of using *in vivo* models of cancer processes in the earliest stages of discovery research. Translationally, we believe the gap in knowledge this study fills is a critical step towards improving treatments for osteosarcoma patients. Many osteosarcoma patients present with undetectable micrometastases, limiting the utility of therapies targeting dissemination and seeding from the primary tumor, as this stage of disease has passed and is largely irrelevant to the patient’s outcome. Established macrometastases mean the cancer cells already have a stronghold in the patient’s lung and through accelerated growth are ready to evolve resistance to therapies. In both scenarios we have likely missed a key therapeutic window of opportunity. Defining intermediate epigenetic states and upstream regulators required for cells to successfully colonize the secondary organ allows us to target metastasis when it is at its most vulnerable. Since patients are likely to possess metastatic cells at different stages of progression simultaneously, a combination therapy tailored to address each distinct biology may be optimal. Although our work identifies reproducible epigenetic changes during osteosarcoma lung colonization, additional studies are needed to assess if the same regulators play roles in other lung metastatic cancers, or even other cancers of the same lineage.

## Methods

### Cell culture

The human osteosarcoma cell lines MG63.3-GFP, MG63.3-GFP-Cas9i, and 143b-HOS-GFP were obtained, generated, and cultured as previously described ^23^. Routine testing for mycoplasma was performed using a custom PCR-based assay.

### Mouse studies

All mouse experiments were performed using NOD scid gamma mice purchased from the Case Western Reserve University (CWRU) Athymic Animal & Preclinical Therapeutics core facility. Mice were housed in ultraclean facilities in accordance to protocols approved by the CWRU Institutional Animal Care and Use Committee. Experiments were designed to minimize mouse use, while optimizing statistical power, based on our extensive experience with xenograft models of metastasis. Mice were standardized for sex and age for each individual experiment. Since no subjective measurements were used for any of the described experiments, researchers were not blinded to group assignments.

### Ex vivo lung metastasis assay

The *ex vivo* pulmonary metastasis assay was performed as previously described ^7,28^. In brief, 1×10^6^ cells were injected into 10-12 week old female NOD scid gamma mice through the lateral tail vein. Mice were immediately sacrificed post-injection. The lungs of the mice were insufflated with a 50% medium/agarose solution. The lungs were then resected and placed in ice cold 1XPBS for 30 minutes to allow the medium/agarose solution to polymerize. After 30 minutes, individual lobes of the lung were cut into transverse sections and placed on Gelfoam (Pfizer, catalog 00300090315085) that had been equilibrated in medium in 6-well plates for 24 hours at 37°C. Sections were then cultured *ex vivo* for 21 days at 37°C and 5% CO_2_. Each section was removed from the Gelfoam and imaged in a new cell culture plate at the timepoints listed in each experiment to assess relative tumor burden by GFP+ area. Fluorescent imaging was performed on the Operetta High Content Imaging System (PerkinElmer), and quantification of tumor burden was done via the Acapella Image Analysis software (PerkinElmer). For the KLF4 knockout experiment, three mice were used per condition, with 6 lung sections cultured per mouse.

### *In vitro* cell viability assay

To measure the effect of *KLF4* knockout on cell viability *in vitro*, 500 *KLF4* knockout cells or non-target control cells were plated in 96-well plates in 90 uL of medium, with six wells seeded per condition. In total, five 96-well plates were seeded to allow measuring of viability at days 0, 1, 3, 5, and 7. After seeding, cells were allowed to adhere overnight before measuring viability at day 0 using PrestoBlue reagent (ThermoFisher, A13261). To do this, 10 uL of PrestoBlue was added to each well and allowed to incubate at 37°C for 1 hour. Fluorescence was then measured on a Synergy Neo2 plate reader (BioTek). For cells cultured past day 0, medium with 1 ug/mL doxycycline was added on day 0. Viability was measured at each timepoint as described above.

### Western blot

Cells were lysed at 4°C in RIPA buffer supplemented with protease inhibitors (Roche, 4693159001). Protein concentrations were measured using a BCA Assay Kit (Thermo Fisher, 23225), and 30 μg of total protein was resolved on precast 4%–12% Bis-Tris gels (Invitrogen, NP0321BOX) and transferred to PVDF membranes (Bio-Rad, 1704157) using the trans-blot turbo transfer system (Bio-Rad, 1704150). The membranes were blocked with 5% dry milk in PBS supplemented with 0.2% Tween-20 (PBS-T) at room temperature for 1 hour and then incubated overnight at 4°C with the following primary antibodies: KLF4 (R&D, AF3640) and cyclophilin B (abcam, catalog ab16045). Chemiluminescent detection was performed with HRP-conjugated secondary antibodies (Santa Cruz Biotechnology Inc., catalog sc-2004; Cell Signaling Technology, catalog 7076S) and developed using Genemate Blue Ultra-Autoradiography film (VWR, 490001-930).

### CRISPR screen and individual inducible knockout cell line generation

The custom CRISPR-screen library was generated as previously described ^36^. Three hundred and sixty sgRNAs targeting the TF targets were pulled from the genome-wide Brunello library ^37^. In addition, 25 sgRNAs without recognition sites in the human genome were included as negative controls. All sgRNAs were synthesized (CustomArray) and cloned into pLV-U6-gRNA-UbC-DsRed-P2A-Bsr; a gift from Charles Gersbach (Addgene, plasmid 83919). Lentivirus was produced with LentiX Packaging Single Shots (Clontech, 631278) according to the manufacturer’s protocol and was used to transduce MG63.3-GFP-Cas9i cells at an MOI of ∼0.3. After selection of the successfully transduced cells with 5 ug/mL blasticidin, screening pools were expanded and used for the *in vitro* and *in vivo* screens. All single knockout cell lines were generated as described above, with the exception that single guides were cloned into pLV-U6-gRNA-UbC-DsRed-P2A-Bsr according to the Broad institute protocol. Knockout induction was performed *in vitro* by exposing cells to 1 ug/mL doxycycline-hyclate (Cayman, 14422).

### sgRNA sequences

sgRNA sequences were as follows: *KLF4*, forward, KO: CACCGAGCGATACTCACGTTATTCG; *KLF4*, reverse, KO: AAACCGAATAACGTGAGTATCGCTC; nontargeting control-1, forward, KO: CACCGAAAAAGCTTCCGCCTGATGG; and nontargeting control-1, reverse, KO: AAACCCATCAGGCGGAAGCTTTTTC.

### In vitro CRISPR drop out screen

For the *in vitro* screen, 5×10^5^ cells were seeded in triplicate in T75 flasks. Cells were maintained for 21 days in the presence of 1 ug/mL doxycycline hyclate, with 5×10^5^ cells reseeded at each passage to maintain 500x library coverage. At the end of the 21 days, 5×10^5^ cells were collected for genomic DNA extraction using red blood cell lysis solution (Qiagen, 28606).

### In vivo CRISPR drop out screen

Fifteen mice were placed on water with 2 mg/mL doxycycline hyclate for 4 days prior to the beginning of the experiment. In parallel, 1×10^6^ cells were seeded in triplicate and cultured in the presence of 1 ug/mL doxycycline hyclate to induce Cas9 expression. Cells were then expanded *in vitro* for 4 days prior to injection. Three groups of 5 mice (15 total) were seeded with 1×10^6^ cells each. Mice were maintained on 2 mg/mL doxycycline for the duration of the experiment. After 21 days of *in vivo* growth, cells were isolated from the lungs of individual mice. In brief, lungs were removed from post-mortem mice and dissociated using the Miltenyi Human Tumor Dissociation kit (Miltenyi, 130-095-929) and GentleMACS dissociator (Miltenyi, 130-093-235) according to the manufacturer’s protocol for medium tumors. To enhance the purity of human cells within the final cell suspension, mouse cells were removed through negative selection using the Miltenyi Mouse Cell Depletion Kit (Miltenyi, 130-104-694). Genomic DNA was then extracted from the purified population of human cells with red blood cell lysis solution and used to prepare sequencing libraries as previously described ^36^. In brief, 2000 ug of genomic DNA was used as a template for PCR, split across 8 separate reactions for each sample. Phusion high-fidelity master mix was used for all PCR reactions. Libraries were then purified using PCR Clean DX beads (Aline Biosciences), pooled, and paired-end sequenced on a MiSeq (Illumina) using custom read and index primers. Although each group of 5 mice was analyzed as a single replicate to obtain a screening coverage of 500x, individual libraries were generated for each mouse.

Analysis was performed using the web-based CRISPRCloud2 ^38^. Dropout for both the *in vivo* and *in vitro* screens were calculated compared to an input day 0 timepoint. For the *in vivo* screen, all 5 mice from each of the three groups were assigned as a single replicate. Hits for each screen were called based on a false-discovery rate < 0.05, and a log2(fold-change) < -1.

### Single cell ATAC-seq

MG63.3 cells were isolated and processed according to the Nuclei Isolation for Single Cell ATAC Sequencing Demonstrated Protocol (10X Genomics, CG000169) with the following modifications: 1% BSA in PBS was used for the washes, and cells were lysed for 3 minutes on ice. Sequencing libraries were created using the Chromium Single Cell ATAC Reagent Kits User Guide (10X Genomic, CG000168). Briefly, 5,000 nuclei were targeted and tagmented in bulk. Nuclei were then portioned into Gel Beads-in-emulsion (GEMs) with a unique cell barcode per single nucleus. GEMs were amplified first in a linear amplification PCR, after which GEMs were broken and PCR was used to add a sample index and Illumina sequencing handles (P5/P7). Libraries were sequenced at the University of Chicago Genomics Facility on an Ilumina HiSeq 4000 with paired-end 100bp reads.

Single cell ATAC-seq data were aligned and processed using Cell Ranger ATAC v1.2 with the hg19 reference genome. The peak cell matrix and fragment file were further processed using Seurat v4.0.1 to create a Seurat object of the class ChromatinAssay ^39^. Using Seurat v4.0.1, latent semantic indexing (LSI) was used to perform normalization, feature selection, and linear dimensional reduction. UMAP reduction was performed using LSI on dimensions 2 through 30 and cells were plotted in UMAP space.

To assess promoter accessibility of marker TFs for each cluster at the single-cell level, Seurat’s FeaturePlot function was used. The promoter for each gene was defined as the genomic region 1kb upstream and downstream from the transcription start site obtained from the UCSC Genome Browser. For visualization, the accessibility score at the promoter of each gene was projected onto the single cell UMAP space.

To determine the distribution of motif enrichment for each marker TF, chromvar was used. First, the motif position frequency matrix was created using motif information from the JASPAR database (JASPAR2020 matrix for species 9606, homo sapiens). Next, chromvar was used to calculate a motif score per cell for each TF. Seurat’s FeaturePlot function was then used to visualize the motif enrichment score per cell for each of the cluster defining transcription factors in the UMAP space.

### ATAC-seq

ATAC-seq was performed using the omni-ATAC-seq protocol previously described ^40^. For cells profiled *in vitro*, 2.5×10^4^, 5×10^4^, or 1×10^5^ cells were used. For the *in vivo* conditions, 1×10^5^ cells were used. Prior to preparation of the *in vivo* libraries, human osteosarcoma cells were isolated from mice 1 day or 22 days post-tail vein injection. In brief, lungs were surgically removed from post-mortem mice and dissociated using the Miltenyi Human Tumor Dissociation kit (Miltenyi, 130-095-929) and GentleMACS dissociator (Miltenyi, 130-093-235) according to the manufacturer’s protocol for medium tumors. To enhance the purity of human cells within the final cell suspension, mouse cells were removed through negative selection using the Miltenyi Mouse Cell Depletion Kit (Miltenyi, 130-104-694). For the early condition, the final cell suspensions from 4 mice were combined prior to performing ATAC-seq for each replicate. For the late condition, each replicate was generated from an individual mouse. MG63.3 libraries were sequenced at the CWRU Genomics Core Facility on an Illumina NextSeq (high-output flowcell) with paired-end 75bp reads. 143b-HOS-GFP libraries were sequenced at MedGenome with paired-end 100bp reads.

Reads were aligned to human genome reference hg19 with BWA-MEM ^41^. Although the purity of human cells was enriched experimentally through magnetic sorting, a residual population of mouse cells were still present in the purified population. To discount these cells from downstream analysis, we also aligned each sample to mouse genome reference mm9 and used the R package XenofilteR to selectively remove reads with better alignment to the mouse genome than human ^42^. Duplicate reads were removed from the filtered BAM files. The de-duplicated BAMs were then used to call peaks with Genrich (https://github.com/jsh58/Genrich) on ATAC-seq mode with default settings. Bigwig tracks were generated using deeptools “bamCoverage” with normalization by RPGC.

To partition the landscape of open chromatin identified based on accessibility dynamics, a z-scored matrix was created from the quantile-normalized fpkm values across the universe of ATAC-seq peaks across all timepoints. Peaks with a coefficient of variation < 10% were binned into a pseudo “static” cluster. The remaining dynamic regions were then clustered with k-means clustering.

### RNA-seq

Cells grown *in vitro* and cells remaining after the *in vivo* ATAC-seq were lysed using TRIzol (Invitrogen, 15596026). RNA was subsequently extracted by transferring the aqueous phase from the TRIzol-chloroform extraction to RNeasy columns (Invitrogen, 74104). The rest of the RNA extraction was performed according to the RNeasy manufacturer protocol. Purified RNA was sent to MedGenome for library preparation and sequencing. MG63.3 libraries were prepared using the Takara SMARTer Stranded Total RNA-Seq Kit v2 - Pico Input Mammalian (Takara, 634411) while 143b-HOS-GFP libraries were prepared using the SMART-Seq v4 Ultra Low Input RNA Kit (Takara). All libraries were sequenced paired-end 100/150bp.

Raw RNA-seq reads were aligned to hg19 and mm9 using HISAT2, and XenofilteR was again used to remove reads that aligned better to the mouse genome than the human genome ^42,43^. Transcripts per million were calculated for each gene across all timepoints.

### ChIP-seq

ChIP-seq was performed as previously described using an antibody targeting H3K27ac (Abcam, ab4729) ^44^. In brief, 5 million MG63.3 sgKLF4 or sgNT cells were fixed with methanol free formaldehyde for 10 minutes. Nuclei were extracted and chromatin was sheared for 7 minutes using the Covaris S2 AFA focused ultra sonicator (Duty factor 5%, intensity 4, 200 cycles/burst). Libraries were prepared as previously described ^44,45^. Libraries were sequenced with paired-end, 150bp reads.

### ChIP-Seq data processing

Cutadapt v1.9.1 was used to remove paired-end adapter sequences and discard reads with a length less than 20bp ^46^. FASTQs were aligned to hg19 using BWA-MEM with default parameters in paired-end mode. Output SAM files were converted to binary (BAM) format, sorted, indexed, and PCR duplicates were removed using SAMtools v1.10 ^47^. Peaks were detected with MACS v2.1.2 with the --broad flag set ^48^. DeepTools v3.2.0 was used to generate RPGC-normalized bigWig tracks with 50 bp bin sizes from the final sample BAM files ^49^. BigWigs were visualized on the Integrative Genomics Viewer in order to assess pronounced track irregularities or low signal-to-noise ratio ^50^. CRISPResso2 was used to confirm editing of KLF4 within the sgKLF4 H3K27ac ChIP-seq data ^27^.

### GREAT analysis

Bed files for each cluster were uploaded individually, using whole genome (hg19 assembly) as the background. To assign genomic regions to genes, all regions from each cluster were uploaded. The region-gene associations were performed with the “Basal+extension” setting, and the resultant pairings were downloaded. To determine enriched biological programs for each cluster, only those regions that showed a log2fc > 1 when comparing either *in vivo* time point to *in vitro* were used.

### Motif analysis

Putative cluster-specific transcription factor regulators were identified through differential motif-mining analysis using GimmeMotifs maelstrom ^51^. Peak lists for each cluster were catted together and reformatted according to the two-column input option. Gimme maelstrom was then run with default parameters. Motifs visualized in the heatmap are those that meet a z-score threshold of 6.

### Statistics

Data in figure 5 are displayed as mean +/- SD. Significance values are calculated by student’s t-test. *** < 0.0005, ** < 0.005, * < 0.05, n.s > 0.05.

## Acknowledgements

This research was supported in part by grants from the National Institutes of Health R01CA193677 (P.C.S.), T32GM088088 (W.D.P.), F31CA247266 (W.D.P.), F31CA213965 (I.M.B.), and T32GM007250 (E.S.H.). We thank members of the Scacheri lab and Tesar lab past and present for their conceptual and technical feedback. We thank Dr. Stevephen Hung for helpful discussion regarding interpretation and early stage writing. We also thank Drs. Michelle Longworth, Yan Li, Jacob Scott, and Yogen Saunthararajah for their advice regarding this work. Additional support was provided by the CWRU Genomics Core Facility, the CWRU Small Molecule Drug Development Core Facility, the CWRU School of Medicine Department of Genetics and Genome Sciences, and the CWRU School of Medicine Department of Molecular Medicine.

## Author contributions

W.D.P. and P.C.S. conceived of the project and designed the experiments. Z.J.F. designed and prepared the CRISPR screen library. W.D.P. performed the CRISPR screen experiments and analyzed the data. W.D.P. generated and analyzed the ATAC-seq and RNA-seq data. E.S.H performed and analyzed the scATAC-seq experiments. W.D.P. and I.B. performed and analyzed the *ex vivo* metastasis experiments. C.F.B. performed the ChIP-seq experiments, and K.L. analyzed the data. J.G. and C.P. assisted with experimentation. W.D.P. and P.C.S. wrote the manuscript. All authors edited and approved the final manuscript.

---

**Supp Fig 1:**
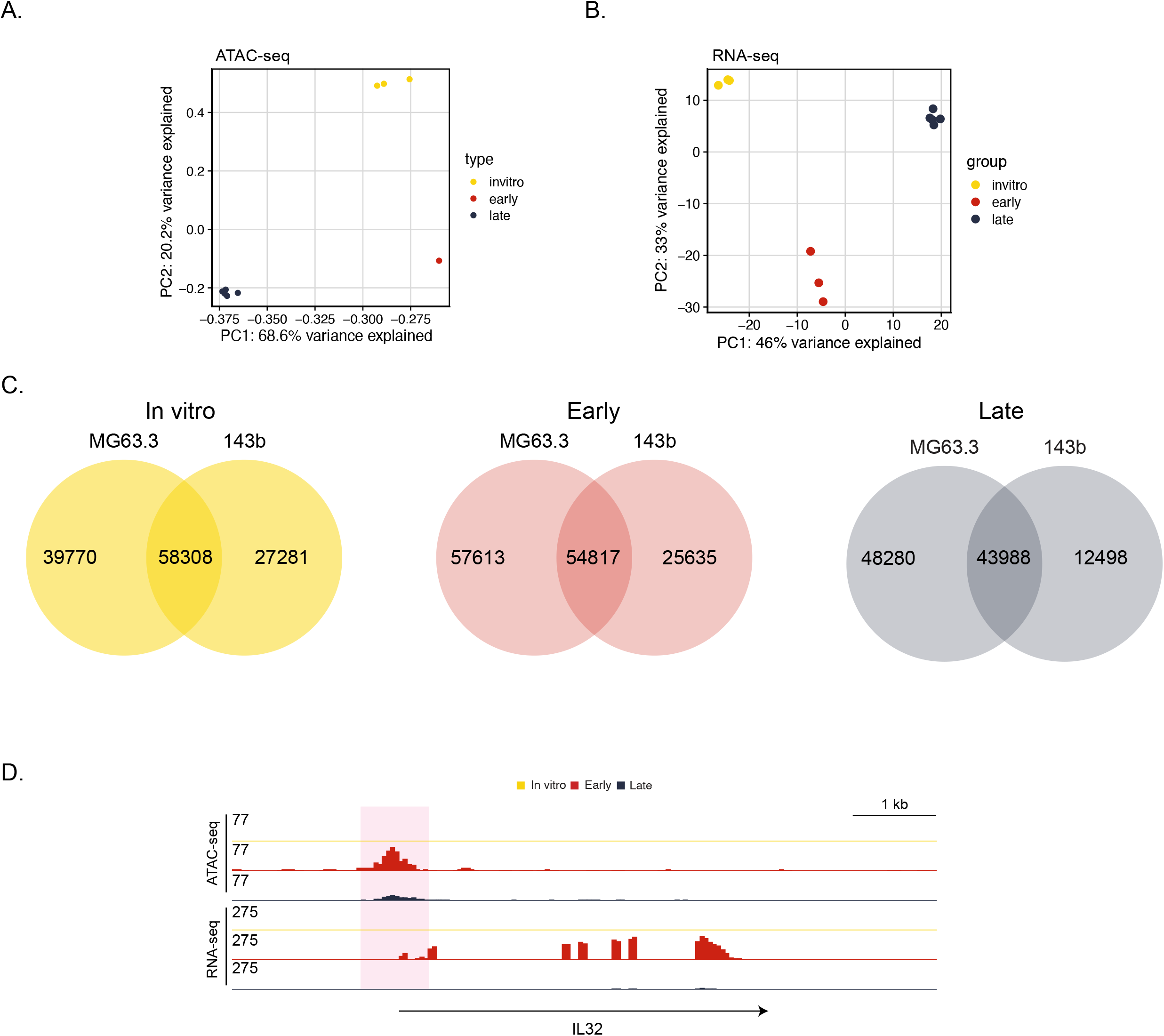
Profiling of chromatin accessibility and gene expression in 143b-HOS during metastasis. **A)** Principal component analysis of open chromatin across the three conditions. **B)** Principal component analysis of gene expression across the three conditions. **C)** Overlap between *in vivo* ATAC-seq data sets from cell lines profiled. Number of peaks in each set are displayed in the venn diagram. **D)** Genome browser view of an early-specific change in chromatin accessbility with corresponding increases in gene expression.

**Supp Fig 2:**
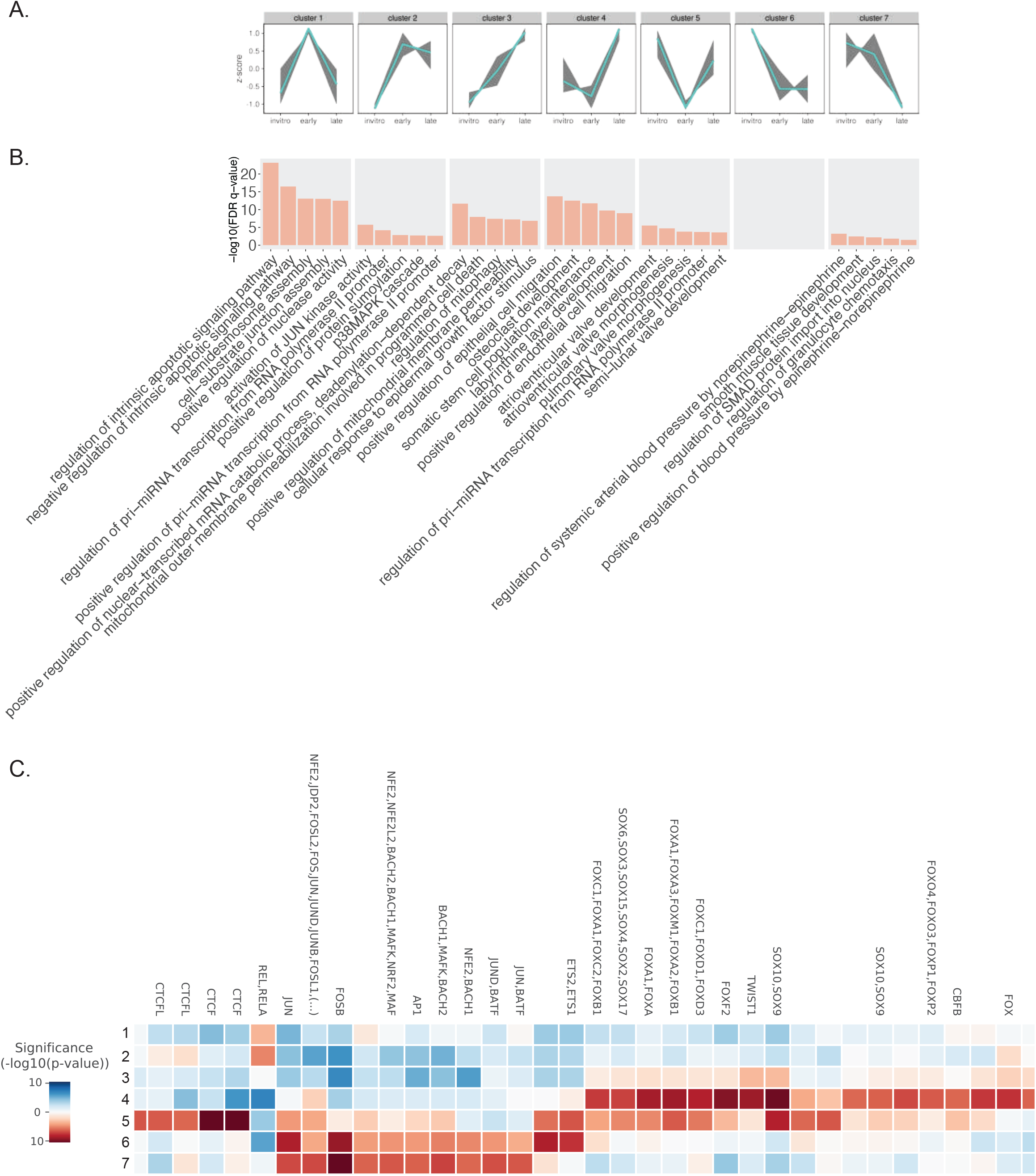
Identification of dynamic clusters of accessible chromatin in 143b-HOS. **A)** Clusters of dynamic accessible regions identified in the 143b cell line based on k-means clustering. The teal line in the middle represents the mean change in accesibility for all peaks within a given cluster. **B)** Peak ontology of each clusters based on GREAT. Top 5 terms for each cluster are shown. **C)** Differential motif enrichment for peaks within each dynamic cluster.

**Supp Fig 3:**
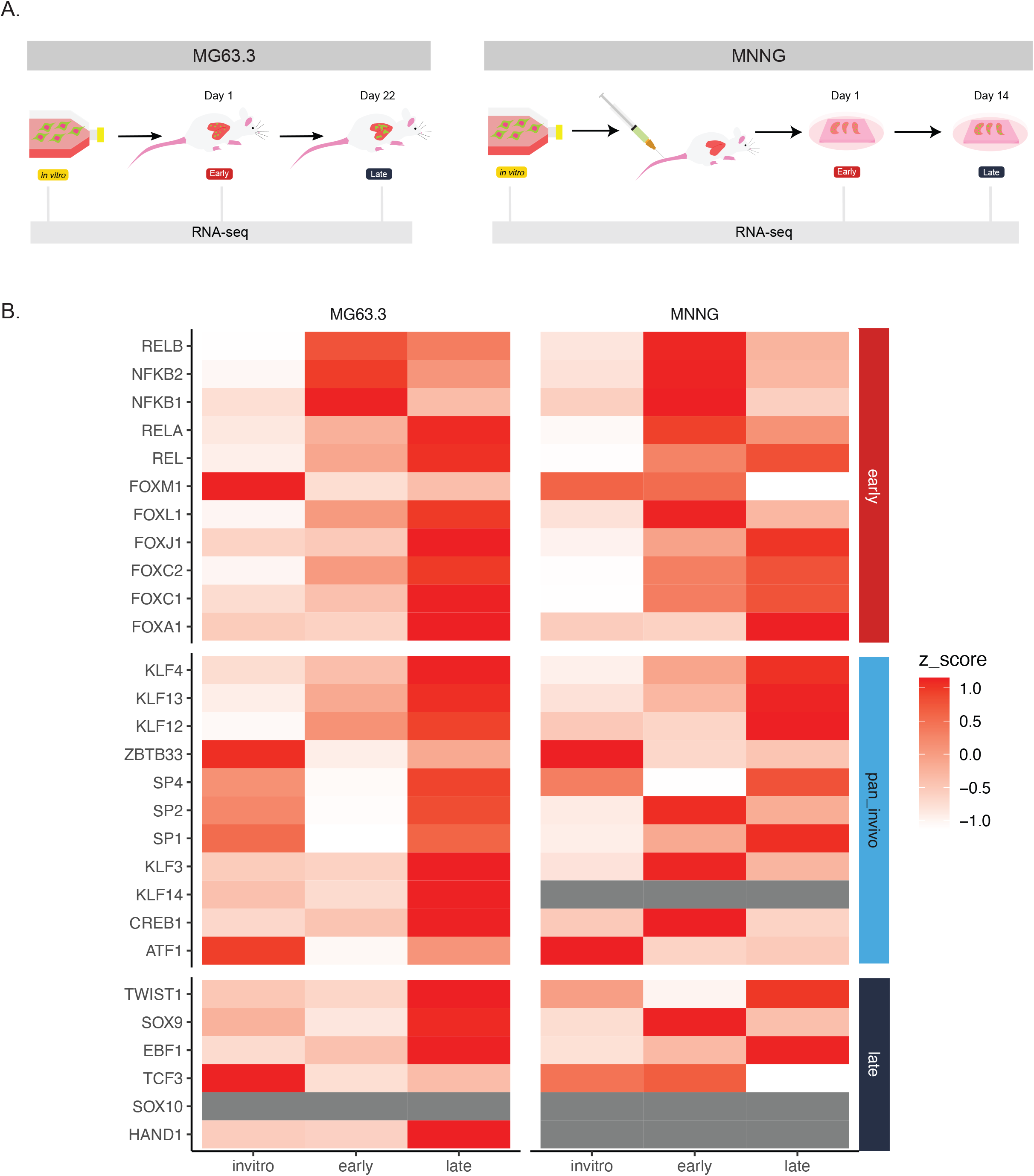
Cluster regulators show similar expression dynamics across multiple metastatic osteosarcoma cell lines. **A)** Schematic comparing RNA-seq profiling experiments of MG63.3 and MNNG cell lines. MG63.3 early and late conditions were isolated from a fully *in vivo* model of metastasis, whereas MNNG cells were isolated from the *ex vivo* PuMA model. **B)** Row-normalized heatmap displaying trends in expression for all putative cluster-regulating TFs in MG63.3 and MNNG.

**Supp Fig 4:**
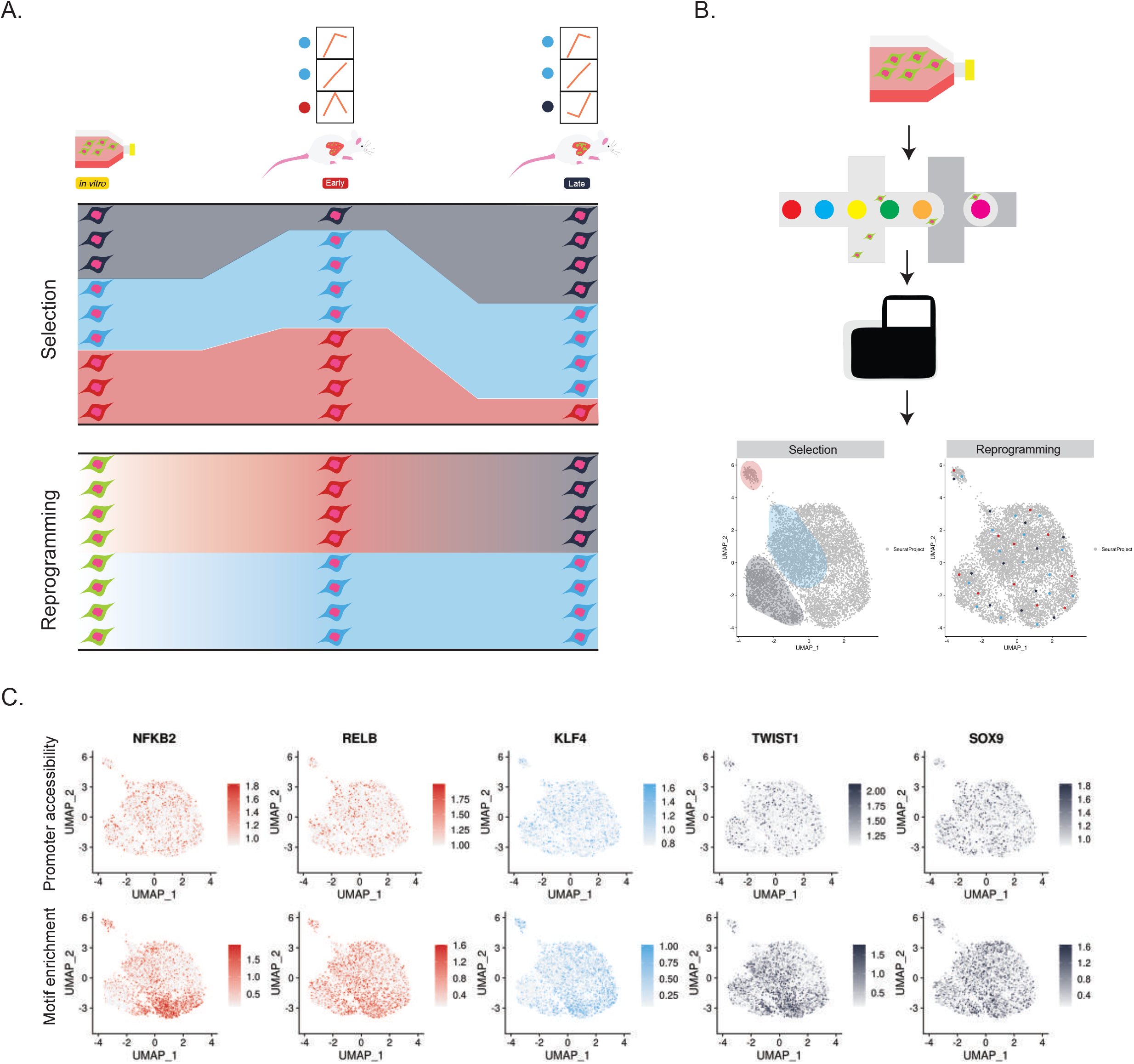
Dynamic clusters of accessible chromatin are not due to subclonal selection *in vivo*. **A)** Schematic of potential causes of dynamic ATAC-seq peaks observed *in vivo*. **B)** Diagram illustrating single cell-ATAC seq experiment and expected distribution of marker transcription factors depending on selection or reprogramming. **C)** Single cell promoter accessibility and motif enrichment for putative dynamic transcription factors.

**Supp Fig 5:**
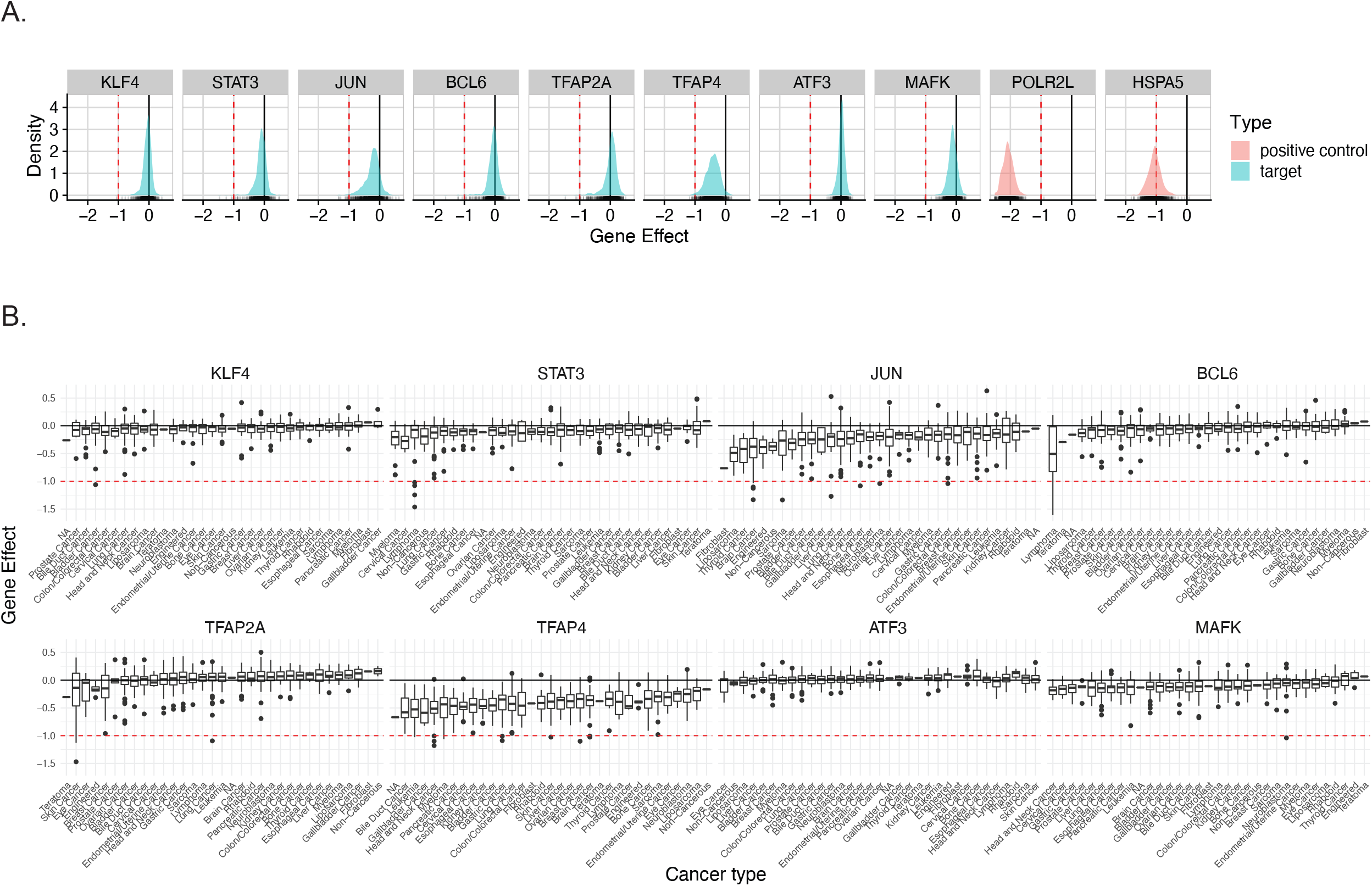
DepMap essentiality data demonstrates specificity of metastasis-dependency genes. **A)** Distribution of gene effect for each *in vivo* hit across all cell lines included in the DepMap database. Genes with an effect less than -1 are dependency genes. **B)** Boxplots highlighting lineage-specific gene effect for each in vivo hit.

**Supp Fig 6:**
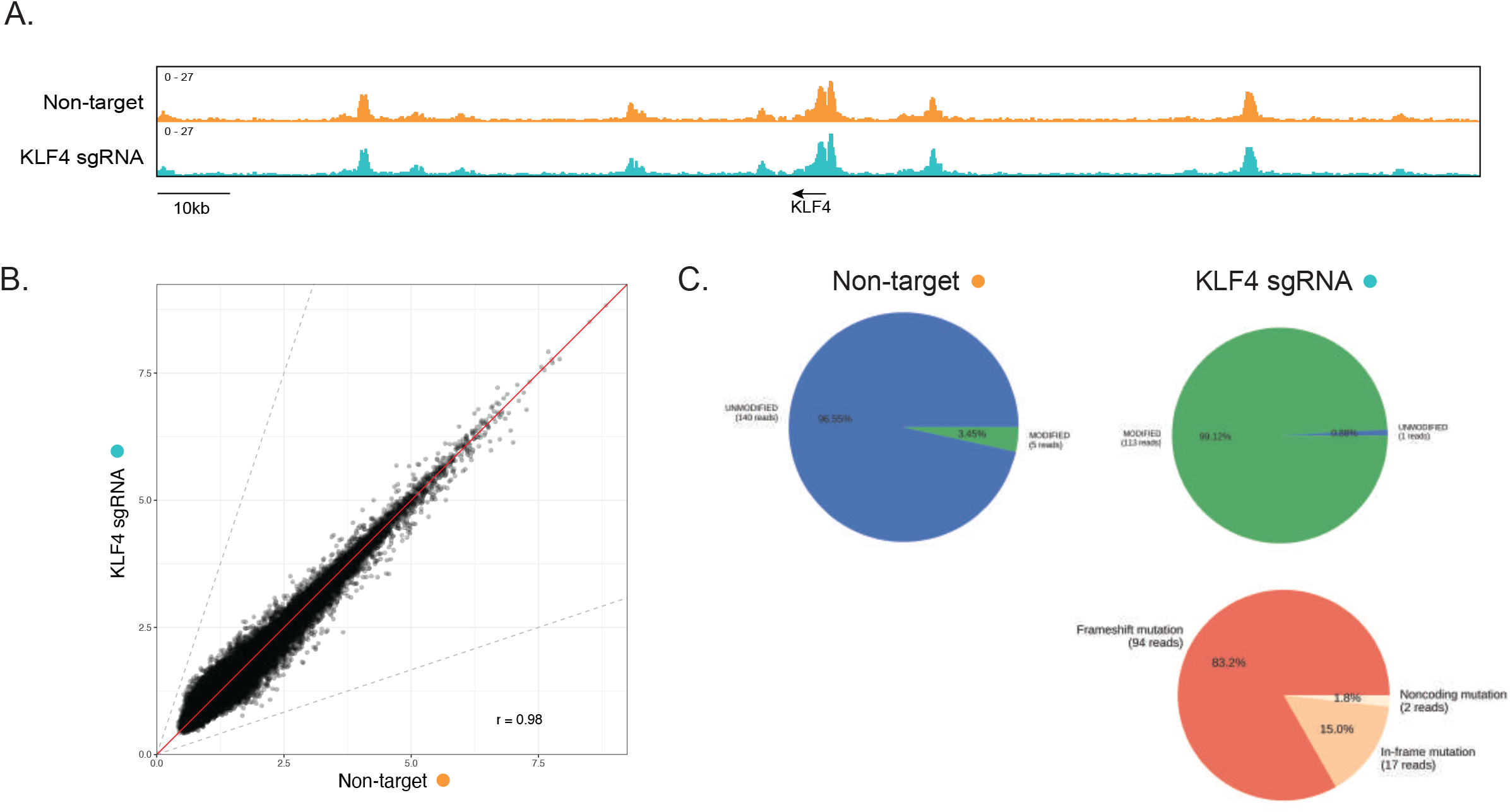
KLF4 knockout does not affect the *in vitro* enhancer landscape of MG63.3 cells. **A)** Genome browser view of H3K27ac signal at the KLF4 locus KLF4 knockout cells and non-target control transduced cells. **B)** Comparison of genome-wide H3K27ac ChIP-seq signal (RPKM) between KLF4 knockout cells and non-target control transduced cells. **C)** Summary of *KLF4* editing for both cell lines based on H3K27ac ChIP-seq reads.

